# Validated Semantic Scoring Improves Filtering of Generic Concepts from Knowledge Graphs for Drug Repurposing Queries

**DOI:** 10.64898/2026.09.29.755063

**Authors:** Frank Hodges, Adilbek Bazarkulov, David Koslicki, Jared C. Roach, Stephen A. Ramsey

## Abstract

When an ontology-based biomedical knowledge graph (a KG, containing information about genes, pathways, phenotypes, and molecular interactions) is analyzed using a computational reasoning system, the system can return unwanted generic concepts. Such generic concepts, in the context of a query about potential drug treatments for a disease, might include ‘pharmaceutical preparations’ or ‘antibiotics’. Answers can be filtered based on hand-curated block-lists or ontology Information Content (IC), but to our knowledge, (1) neither filtering method has been validated against expert judgments of generality, and (2) IC is not available for the majority of nodes in the representative KG used in this work. In this study, three domain experts rated representative lists of KG concepts for generality. We built a semantic precision score (SPS) from a concept’s name and description alone, using a biomedical language model (BioLORD). For the purpose of filtering generic answers from biomedical query results, we compared (using queries for 200 diseases) the SPS against a curated concept block-list and two IC thresholds. IC agreed moderately with the experts (Spearman’s *ρ* = 0.59), but correlations were negligible or negative within IC bands. The SPS was well-correlated with the experts (*ρ* = 0.76) globally and within SPS bands, and remained informative on concepts without IC (*ρ* = 0.59). Hard filtering of query results using the SPS removed 1,502 of 33,161 query answers, nearly three times the block-list’s 513, while removing fewer of the query-specific DrugCentral indications. Hard filtering by IC removed either few answers or many approved drugs. We additionally compared the filtering methods’ performance on an independent evaluation set of queries for 200 diseases. The SPS filter removed 5,135 of 91,203 answers compared with the block-list’s 1,419 and lost none of 1,296 indications compared with 17. Further, SPS filtering removed approved drugs no more often from the half of DrugCentral not used in training than from the half used in training. Semantic meaning captures expert judgments of specificity more faithfully than ontology position, and semantic scoring can replace IC and manually maintained block-lists for filtering drug-repurposing results.

## 1 Introduction

Biomedical knowledge graphs bring together information about concepts of various types, including genes, phenotypes, pathways, diseases, and drugs. Such graphs enable computational reasoning for diverse applications, such as explaining gene-phenotype relationships, analyzing data from genomics studies, or identifying potential drug-repurposing candidates. In a knowledge graph, each concept is represented by a node, and a relationship between two concepts is represented by an edge. Biomedical knowledge graph relationships can come from various sources, including curated knowledge bases, formal ontologies, literature-mining, and predictive models. The NCATS Biomedical Data Translator (‘Translator’) reasoning system [Biomedical Data Translator Consortium, 2019, Fecho et al., 2022] includes, and primarily uses, one such graph, the Translator tier-0 knowledge graph [Biomedical Data Translator Consortium, 2026b]. Within the Translator system, the analytical agent ARAX [Glen et al., 2023], which was co-developed by the authors of this work, answers queries by returning candidate entities, such as chemical substances or drugs, that may be therapeutically beneficial for a disease specified in the query context; we refer to such queries as treats queries. In the Translator system, `treats` query results frequently contain broad class-level concepts such as ‘Pharmaceutical Preparations,’ ‘Antibiotics,’ and ‘Immunomodulators,’ which we collectively call *generic* concepts. As query results, generic concepts may be formally valid by virtue of being connected to the query’s fixed concept (such as the disease) in the knowledge graph, but are of lower value than results based on specific concepts such as individual drugs.

Several strategies have previously been described for reducing generic concepts’ appearance in, and/or influence on, graph query results. The SemMedDB database of literature-mined knowledge includes a curated list of generic concepts [Kilicoglu et al., 2012], which Zhang et al. [2021] used to filter out relations involving generic concepts. Filtering is also used contextually based on semantic type; Zhang et al. excluded certain concept node types and edge relationship types that were deemed irrelevant to the query context. Graph topological heuristics have also been employed to reduce generic results: given that generic concepts are often highly connected in the knowledge graph, Zhang et al. and Preiss and Stevenson [2017] filtered out highly connected nodes, and Himmelstein et al. [2017] down-weighted graph paths through highly connected nodes. Other graph topological heuristics are based on a concept’s position in a formal ontology: Resnik [1995] proposed Information Content (IC), which measures a concept’s specificity based on its position in the ontology’s class hierarchy; a concept with many descendants in the hierarchy has low IC and a leaf concept has an IC value that is at least as large as the maximum IC of its ancestors [Resnik, 1995, Seco et al., 2004]. IC has since been adopted into the human phenotype ontology [Köhler et al., 2009], the gene ontology [Supek et al., 2011], the Ubergraph integrated biomedical ontology [Balhoff et al., 2022], the Babel concept database [Biomedical Data Translator Consortium, 2026a], and (via Babel) the Translator tier-0 graph. IC can be used to reduce the influence of generic concepts in similarity-search or representative-selection tasks [Köhler et al., 2009, Supek et al., 2011].

ARAX currently suppresses generic answers using a curated block-list of concepts (numbering 4,193 at the time of writing). The Translator tier-0 knowledge graph also provides IC values for nodes that have such values available, making IC a potential alternative to manual filtering of query results. However, using IC for this purpose assumes that (1) for concept nodes that may appear in query results, IC is defined (which depends on the node being in an ontology for which IC values are available); and (2) the concept’s position within its ontology reflects absolute semantic specificity. Both of these assumptions are false: (1) in the

Translator graph, IC is defined for only 34% of nodes and is sparse or absent for several groups of concept nodes (of specific semantic type) that are important in query results; and (2) an ontology node with no children in the class hierarchy may still denote a broad class, and a node with many children may denote one specific entity. These two problems motivate (1) creating an improved specificity metric that can be computed for any node based on its textual semantic information (which is always available) rather than its class hierarchy position (which is often not available) and (2) evaluating both the new metric and IC directly against human judgment, which to our knowledge has not been done for topology-based filtering.

This work addresses three questions: **Q1**, how well does IC track expert judgments of specificity? **Q2**, can a new score, which we call the SPS, derived from a concept’s human-readable name and free-text description, better capture those judgments and extend to concepts without IC? **Q3**, for a drug recommendation task using an evaluation set of diseases with known ground-truth answers, from the standpoint of answer correctness and completeness, how does filtering with the SPS compare with the hand-curated block-list and with using IC thresholds? Q1 and Q2 test whether the score measures the intended construct, and Q3 evaluates its behavior in the downstream task for which generic-node filtering is needed. We report empirical findings related to each question. First, we evaluated how well IC reflects expert judgments of concept specificity using two lists of concepts that we developed. Second, we defined and evaluated a concept specificity model, the SPS, that needs only a concept’s text. The SPS combines a generality direction learned without expert labels with a biomedical language model trained on labeled examples, evaluated once against the held-out expert panel. Third, in the context of using ARAX to run `treats` queries for 400 diseases (an initial set of 200 whose query results also informed selection of the SPS, and an independent set of 200 used to evaluate the fixed model) with known ground-truth correct answers, we evaluated three different generic concept filters: thresholding the SPS, thresholding IC, and using the ARAX block-list. Our evaluation included measuring, for each disease, the number of approved drugs that were wrongly removed and the count (among the top ten results for the disease) of drugs that are indicated for the disease.

## 2 Materials and Methods

In this section, we describe our approach for scoring the semantic precision of concept nodes in biomedical knowledge graphs, and for comparing the SPS to alternative generic concept filters based on IC or the ARAX block-list. Full technical details are available in the Supplementary Information.

### 2.1 Graph and Information Content

We based our analysis on a representative up-to-date/comprehensive knowledge graph built for biomedical reasoning: the Translator tier-0 graph, version 2026-06-21 [Biomedical Data Translator Consortium, 2026b], which has 1.75M nodes and 28.7M edges. The properties of each node include a human-readable name, a semantic type ‘category’ from the Biolink semantic model [Unni et al., 2022], and, for 34.5% of nodes, a description. In the tier-0 knowledge graph, an IC score is available for only 34.3% of nodes, mostly corresponding to specific node categories that have comprehensive ontologies (Fig. 1) such as BiologicalProcess (93%) and Disease (84%); for `treats` queries, the low IC coverage of nodes in the node categories Drug (0%) and ChemicalEntity (22%) is particularly problematic for genericness filtering.

**Figure 1:**
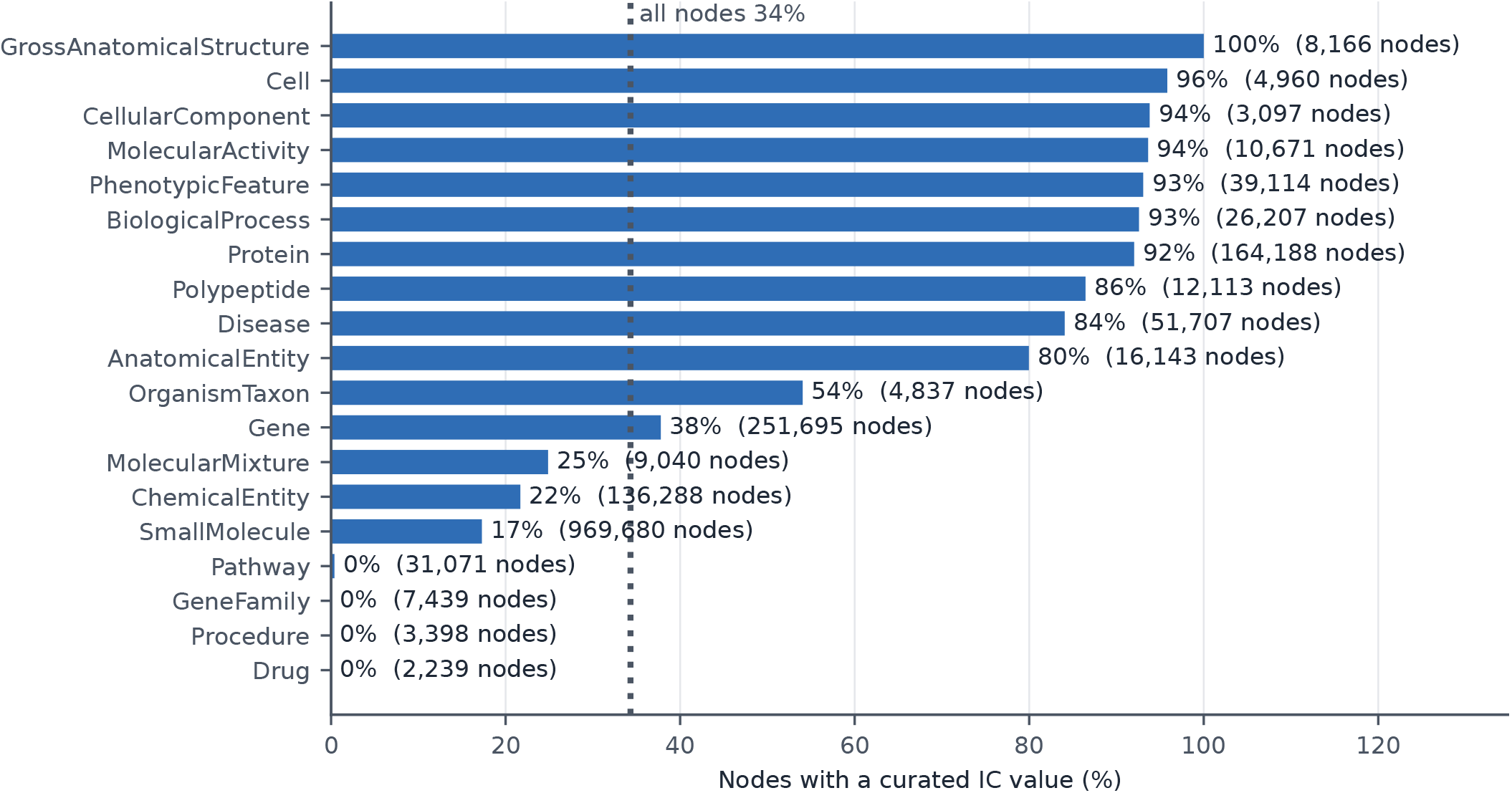
Share of nodes with a curated IC value, by Biolink category.

### 2.2 Expert panel for rating concept nodes’ specificity

Three domain experts (JCR, SAR, and DK; all co-authors on this work) independently annotated concepts for semantic specificity through per-rater forms in randomized order. The domain experts had no involvement in training, tuning, selection, or evaluation of the machine-learning-based specificity models developed in this work. For each concept, the raters were provided the concept’s name, description, and category, but not IC, model scores, or each other’s ratings. For each concept, each rater assigned an integer from 1 (generic, too general to be a useful standalone answer to a biomedical query) to 5 (specific, a concrete useful answer), judging specificity only based on the provided node properties, and without accessing additional online resources.

Each expert assigned ratings to all concepts on two concept lists that were generated by a stratified sampling technique as explained below. List A has 100 nodes with IC, 20 per IC band (0 to 50, 50 to 66, 66 to 88, 88 to 95, 95 to 100), at most three per Biolink category per band, with a guard used to ensure that no near-duplicate concepts were on the list. List A was purposefully band-balanced, so pooled statistics describe a balanced sample and not the graph, where the top band dominates. List B has 50 nodes without IC, drawn uniformly within category and weighted toward the zero-IC categories where `treats` query answers reside (Drug, ChemicalEntity, SmallMolecule, Pathway, GeneFamily, Procedure among them). Only 10% of List B items have a description. Both lists and the SPS model (see Sec. 2.5) were frozen before rating. The consensus scale (‘panel’) is the mean rating.

### 2.3 The first-stage embedding model and its ‘genericness’ score

As a first-stage measure of concept generality, we developed a ‘genericness’ score for concept nodes, based only on two textual properties of a concept node (the human-readable name and the free-text description), using a large language model to map the textual information into a vector in an embedding space and then using a linear model to compute the genericness score. For the embedding language model, we used SapBERT [Liu et al., 2021], which maps a node’s textual information to a real vector of length 768. The linear model’s feature space was the SapBERT embedding space, and the model was fitted to a large representative list of pairs of parent/child concept nodes (in the sense that the child node is a subclass of the parent node) that we created using a text-based approach. For each pair, we took the difference of the parent and child embeddings and fitted a unit-norm vector *w* without intercept by L2-regularized logistic regression, using both orientations of every pair, so that *w ·* (*e*_parent_ *− e*_child_) *>* 0, i.e., the parent projects higher than its child. For the text-based approach to generate the representative list of parent-child concept pairs, we implemented a lexical rule that pairs two names when the parent name ends the child name, as in ‘transport’ and ‘sodium ion transport.’ A second (definitional) rule takes the genus term from Open Biological and Biomedical Ontology (OBO) definitions, so ‘Any *metabolic process* that… ‘ gives ‘metabolic process’ as the parent. Together, the two rules produced 660,663 parent-child pairs, involving over 93,693 distinct parents. We searched for the unit vector that minimized the concept genericness function using stochastic gradient descent. With the linear model trained, the predicted genericness score for any concept node (whether or not it was in the representative set of parent/child pairs used for model fitting) is the linear model’s prediction for the node’s embedding vector, standardized over all nodes (higher is more general). Importantly, the ‘genericness’ score model was not trained using a ground-truth set of concept nodes labeled ‘generic’ or ‘not generic’; instead, its training was supervised only by a set of parent/child concept node pairs extracted using text-based heuristics. We then incorporated the ‘genericness’ score as a feature in a second-stage model, the SPS (Sec. 2.5), that was trained using a large set of concepts labeled generic/specific.

### 2.4 Labeled data set for training the SPS model

For supervised training of our SPS model, we developed a representative set of 34,915 concept nodes labeled ‘generic’ or ‘specific’, as follows. We started with the ARAX generic concept block-list, which we reduced via filtering and semantic normalization to a set of 1,053 generic concept nodes in the tier-0 graph. Next, we mapped all 1.75M concept nodes in the Translator tier-0 graph (specifically, their text information consisting of the human-readable concept name and a free-text description field) to points in the SapBERT embedding space (Sec. 2.3). Using positive-unlabeled learning, with the ARAX block-list concepts as positives, we trained a gradient-boosted decision tree (GBDT) classifier on ten node features derived from the graph connectivity, its IC, and its embedding (Supplementary Table S1). Negative examples were drawn from unlabeled nodes at a 10:1 ratio of negatives to positives, with removal of sampled negative nodes for which the node name is only the node’s accession number (a ‘bare accession’). The classifier assigned each of the 1.75M nodes a score between 0 and 1 (higher is more likely generic). For the 25,595 nodes with GBDT score exceeding 0.8, we used a locally-hosted large language model (Qwen2.5) to assign the label ‘generic’ or ‘specific’ to each node based on the node’s textual information, in accordance with a four-rule prompt: (1) a class of things is generic; (2) one identifiable thing is specific; (3) breadth is not genericness; and (4) accessions and gene symbols are specific. The Qwen2.5 model labeled 8,980 concepts generic and 16,615 concepts specific. We also explored convergence of LLM-in-the-loop training of the GBDT over multiple iterations (see Supplementary Information).

In addition to the Qwen2.5-labeled concepts, we extracted from DrugCentral [Ursu et al., 2017] a moderate-size (but high-confidence) set of negative (‘specific’) concepts. Specifically, we collected the subject (or head) nodes of DrugCentral `treats` edges in the tier-0 graph (corresponding to 2,330 drug nodes). From these negative examples, we allocated 1,127 for training (Sec. 2.5), and held back 1,203 for validation. Concepts that were also in List A for expert rating (Sec. 2.2) were excluded from the labeled set. This yielded a set of 34,915 labeled nodes (10,004 positives and 24,911 negatives).

### 2.5 The SPS for a concept node’s semantic precision

To develop a highly accurate text-based score for the semantic precision of concept nodes, we required a language embedding model with a deeper semantic understanding of biomedical concepts. For this purpose, we chose BioLORD-2023 [Remy et al., 2024], which is pre-trained for mapping concept names to definitions. As with the SapBERT-based first-stage genericness score model, for the SPS model, we embedded (using BioLORD) a concept node’s textual information (its human-readable name and free-text description). We implemented a linear prediction layer whose 769 inputs are the concatenation of the genericness score (Sec. 2.3) and the 768 BioLORD embedding vector components. We fine-tuned all parameters of the BioLORD encoder jointly with the linear prediction layer (769 weights and a bias, with dropout 0.1 on its input) to minimize the cross-entropy logistic loss *ℓ*(*y, z*) for the training set of examples (Sec. 2.4), treating the output of the linear predictor as the logit *z*. To obtain a SPS that is higher with increasing semantic precision, we defined the SPS as the negative of the linear predictor’s output *z*, i.e., *−z*. Because we weighted the two classes in the loss function to ensure class-balanced contributions to the overall loss, the value *z* is interpretable as the log-likelihood ratio of a concept being generic vs. specific, given its textual information. We used 0 as a filter cutoff for identifying generic concepts: i.e., concepts for which *z >* 0 were treated as generic and filtered out of query results. We jointly trained the linear model and fine-tuned the BioLORD embedding model in three epochs, with 10% of labeled concepts held back to monitor fit, and with a batch size of 48 concepts. We used stochastic gradient descent optimization with learning rates of 2 *×* 10^*−*5^ for the BioLORD model and 10^*−*3^ for the linear model. Training was repeated with three random seeds under the same rule, and the final SPS that we assigned for each concept is the average of the three models’ scores. Lists A and B were scored using the trained model.

For the task of detecting generic concepts, we computed the precision vs. recall trade-off curves for the SPS and for the IC score by sweeping the score thresholds and, for each threshold value, computing (1) the number of high-confidence specific nodes from the DrugCentral list (see Sec. 2.4) that were classified generic (a measure of the false positive error rate); and (2) the proportion of concepts on the ARAX generic concept block-list that were classified as generic (a measure of recall).

### 2.6 Query-level test

To empirically study genericness filters (Q3), we used the open-source ARAX query-answering system (github.com/RTXteam/RTX). From MONDO diseases with at least two DrugCentral indications in the tier-0 graph, we randomly selected 200 diseases. For each disease, we queried ARAX to obtain a ranked list of treatments and then processed the lists offline outside of ARAX. We measured the performance of four generic result filters that exclude results that (1) match the ARAX block-list; (2) have IC below 50; (3) have IC below 80; and (4) have SPS below zero. As a control, we measured performance without generic-result filtering. For the IC filters, results lacking an IC were retained (i.e., assumed specific). Across the 200 disease-specific queries, for each filter type, we calculated four query-result performance measures: ‘Removed’, which we defined as the number of answers removed; ‘Own indications removed’, which we defined as the number of drugs that were removed from a query result that were actually (per DrugCentral) indicated for the disease in the query context (a measure of filtering ‘collateral damage’); ‘DrugCentral drugs removed’, which we defined as the number of DrugCentral drugs that were removed from query results overall (whether or not the drugs were indicated for the disease specified in the query); and ‘Top-ten indications’, which is the number of ground-truth positive drug-disease indications that were listed within the top ten entries in the corresponding disease’s query results.

To control for any bias from overlap of ARAX results from the 200 disease-specific queries and labeled concept nodes that were used in model training/selection, we constructed a replication set of 200 additional diseases. As a check on the expert labeling of generic concepts, we conducted a large language model (LLM) audit (see Generative AI Statement). In brief, we asked two locally-hosted LLMs to rate each of 150 expert-labeled concepts (and 620 concepts appearing in query results) as generic or specific based on the concept’s name and description and based on a judging rubric (Sec. 2.4).

### 2.7 Statistics

For testing bivariate monotonic association, we computed Spearman’s *ρ* and estimated each confidence interval (CI) via 10,000-fold bootstrap resampling. We used Krippendorff’s *α* to measure inter-rater agreement. For binary classification, we measured receiver operating characteristic (ROC) curve area. For replication-set removal counts, we obtained 95% CIs using bootstrap over queries (2,000 resamples).

## 3 Results

### 3.1 Internal consistency of expert panel’s annotations is list-dependent

The expert panel’s (panel-mean) concept genericness ratings had a relatively high inter-rater agreement score of *α* = 0.727 on List A, with pairwise agreements in the range of *ρ* = 0.74 to *ρ* = 0.80, consistent with the expectation that genericness is a well-defined construct on ontology vocabularies. On List B, which contains names only and consists mostly of catalog-style entities (patent compounds, lipid species, product strengths), the agreement was lower, *α* = 0.372, as expected for name-only concepts that are not ontologically rooted.

### 3.2 Q1. IC is a poor predictor of relative degree of concepts’ genericness

On List A concepts, the correlation of IC with the expert panel ratings was relatively low (*ρ* = 0.59, 95% CI 0.46–0.70, Fig. 2A), with significant disagreements both ways (IC greater than panel mean, and IC less than panel mean). Panel means rise across IC bands (2.58, 3.07, 3.78, 4.50, 4.32), but within bands, the correlation is at most 0.34 and is negative at both ends, *−*0.33 in 0 to 50 and *−*0.25 in 95 to 100 (Fig. 3). Given two concepts of similar IC, IC gives no usable order, and the band where it anti-correlates with experts holds 78% of the concept nodes that IC covers. Table 1 shows both example concepts for which IC and the expert panel disagree and example concepts for which they agree. As an example of a type of disagreement, an umbrella term (‘toxin sequestering activity’) that happens to be a class hierarchy leaf in an ontology has a maximal IC because it does not (yet) have any subclasses in the ontology, while experts annotated it as a broad concept. Conversely, richly subclassified specific concepts (‘gamma-aminobutyric acid secretion, neurotransmission,’ L-glutamate) get low IC because salt forms and finer distinctions sit beneath them, while experts judge the concept itself. A cutoff chosen from the panel ratings shows the consequence. With concept nodes for which the expert panel-mean rating was *≤* 2 labeled as generic, the IC cutoff that best balances sensitivity and specificity (IC = 86.3) has sensitivity 1.00 and specificity 0.51, flagging half of the non-generic concepts.

**Table 1:** List A concepts for which IC and the expert panel strongly agree or strongly disagree.

| Concept Name | Biolink Category | IC | Panel |
| --- | --- | --- | --- |
| *IC disagrees with experts:* |  |  |  |
| toxin sequestering activity | MolecularActivity | 100.0 | 2.33 |
| positive regulation of sulfate assimilation | BiologicalProcess | 100.0 | 2.67 |
| GABA secretion, neurotransmission | BiologicalProcess | 42.7 | 4.67 |
| L-glutamate(1-) | ChemicalEntity | 42.8 | 4.00 |
| *IC agrees with experts:* |  |  |  |
| mCode Non-Neoplastic Disease/Finding | PhenotypicFeature | 47.5 | 1.33 |
| tripeptide zwitterion | SmallMolecule | 40.6 | 1.67 |
| *TBL5* | Gene | 100.0 | 5.00 |
| *CMPG2* | Gene | 100.0 | 5.00 |

**Figure 2:**
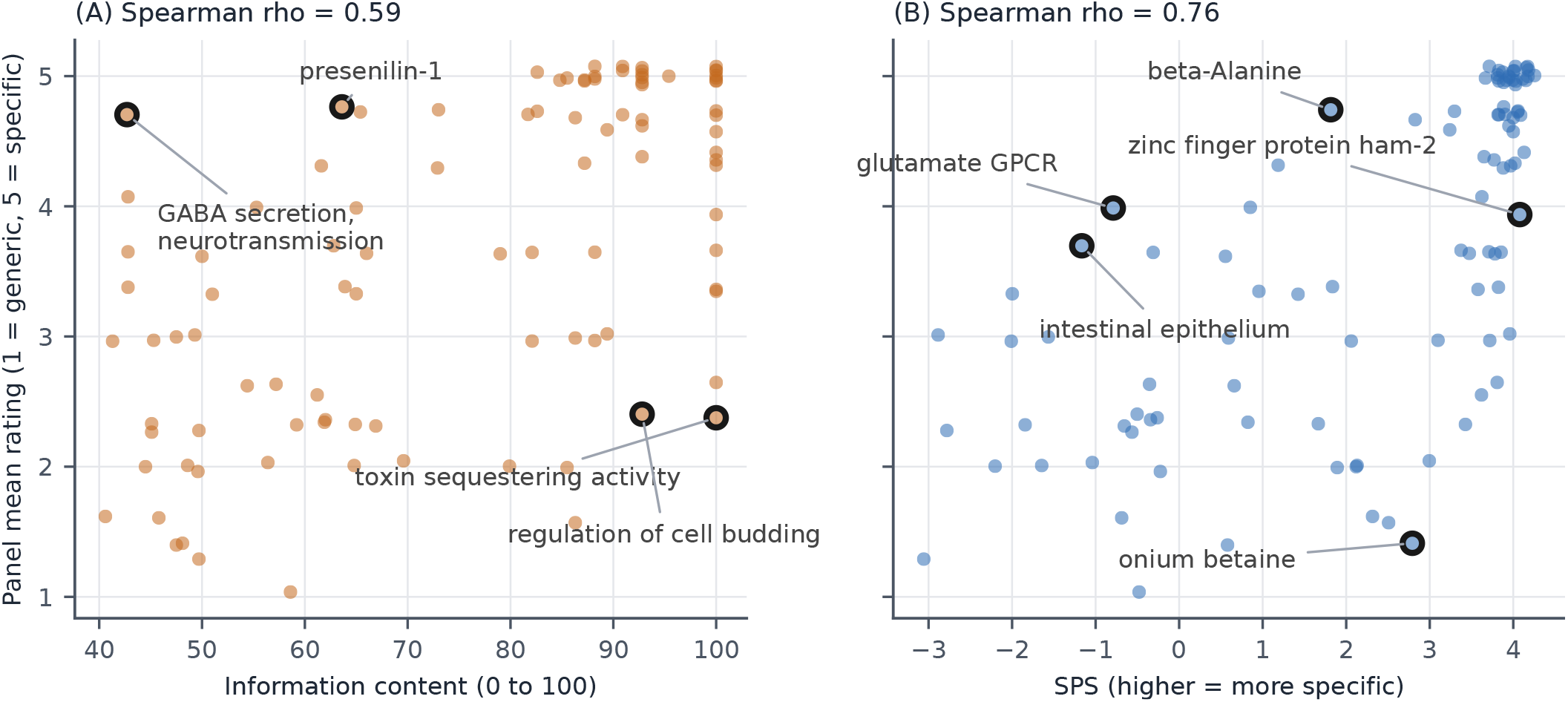
SPS is more consistent with expert ratings of genericness than IC is. Each mark represents a concept node from List A (*n* = 100). Circled points are the largest disagreements (see Table 1). **(A)** IC against the mean expert score. **(B)** The SPS against the panel mean.

**Figure 3:**
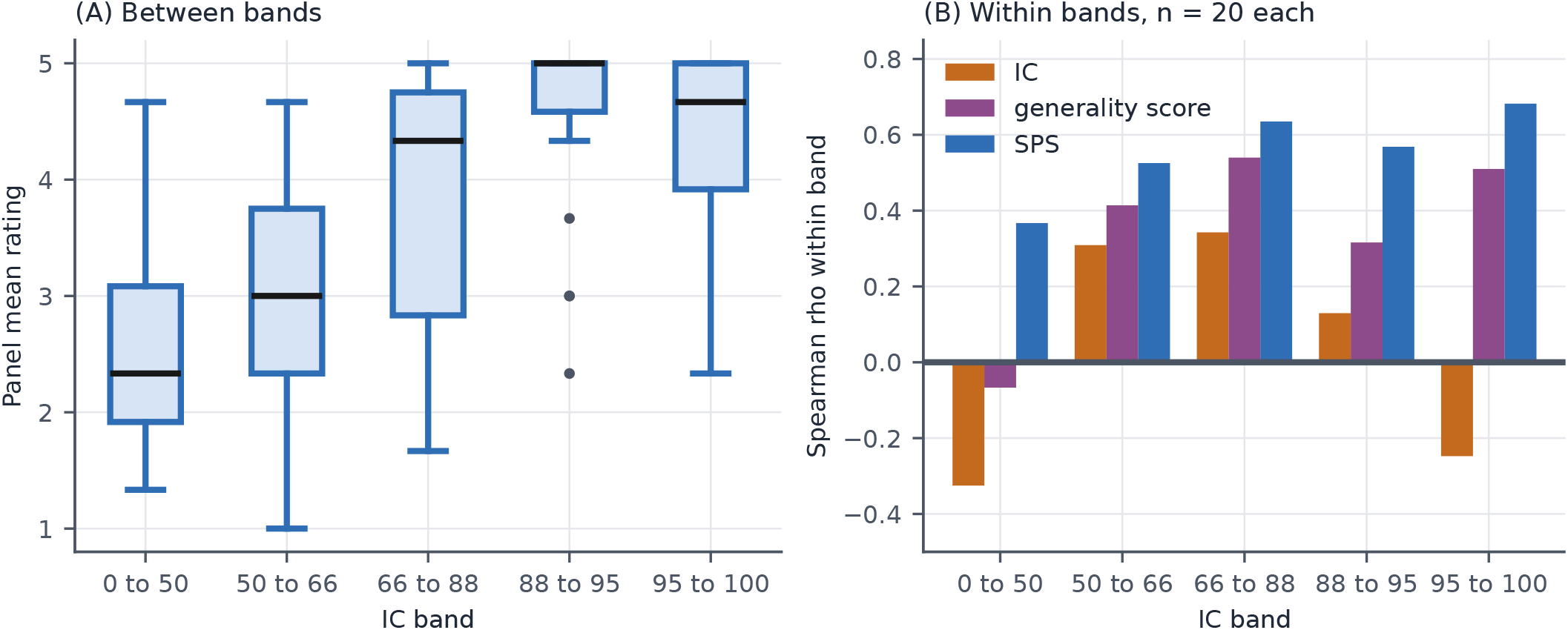
**(A)** Panel mean ratings of concepts grouped by IC band. Black bar denotes median; box denotes interquartile range (IQR); whiskers extend *±*1.5 IQR beyond the IQR. **(B)** Within-IC-band (*n* = 20 per band) Spearman correlations between the expert panel and each of three score types: IC, the generality score, and the SPS.

### 3.3 Q2. The SPS beats IC on the expert-annotated data set

On List A concepts, the SPS was far more consistent with the expert panel (*ρ* = 0.76; 95% CI 0.68–0.81; Figure 2B) than IC, with a correlation that is comparable to the raters’ correlations with one another (with the caveat that a model-to-consensus correlation and a rater-to-rater correlation are not the same quantity, since the consensus averages out rater noise). The SPS’s correlation with expert ratings is consistent across IC bands (Fig. 3B). On the 50 no-IC nodes of List B, the SPS correlates with the expert panel at *ρ* = 0.59 (95% CI 0.35–0.77, Fig. S1).

### 3.4 Q3. On treats queries, SPS filtering is better than IC filtering

Our empirical study of performance of three different ARAX query result post-filters on a set of 200 randomly selected diseases produced a total of 33,161 result answers (a median of 166 answers per disease), with 513 of the answers on the ARAX generic concept block-list. Filtering out answers with SPS *<* 0 removed a significantly higher proportion (4.5%) of generic answers than the IC*<* 50 filter (0.6%), while at the same time removing fewer known-correct answers (SPS removed 2; IC*<* 50 removed 3) and removing fewer known-specific answers as determined by DrugCentral (SPS removed 28; IC*<* 50 removed 91) (Fig. 4A).

**Figure 4:**
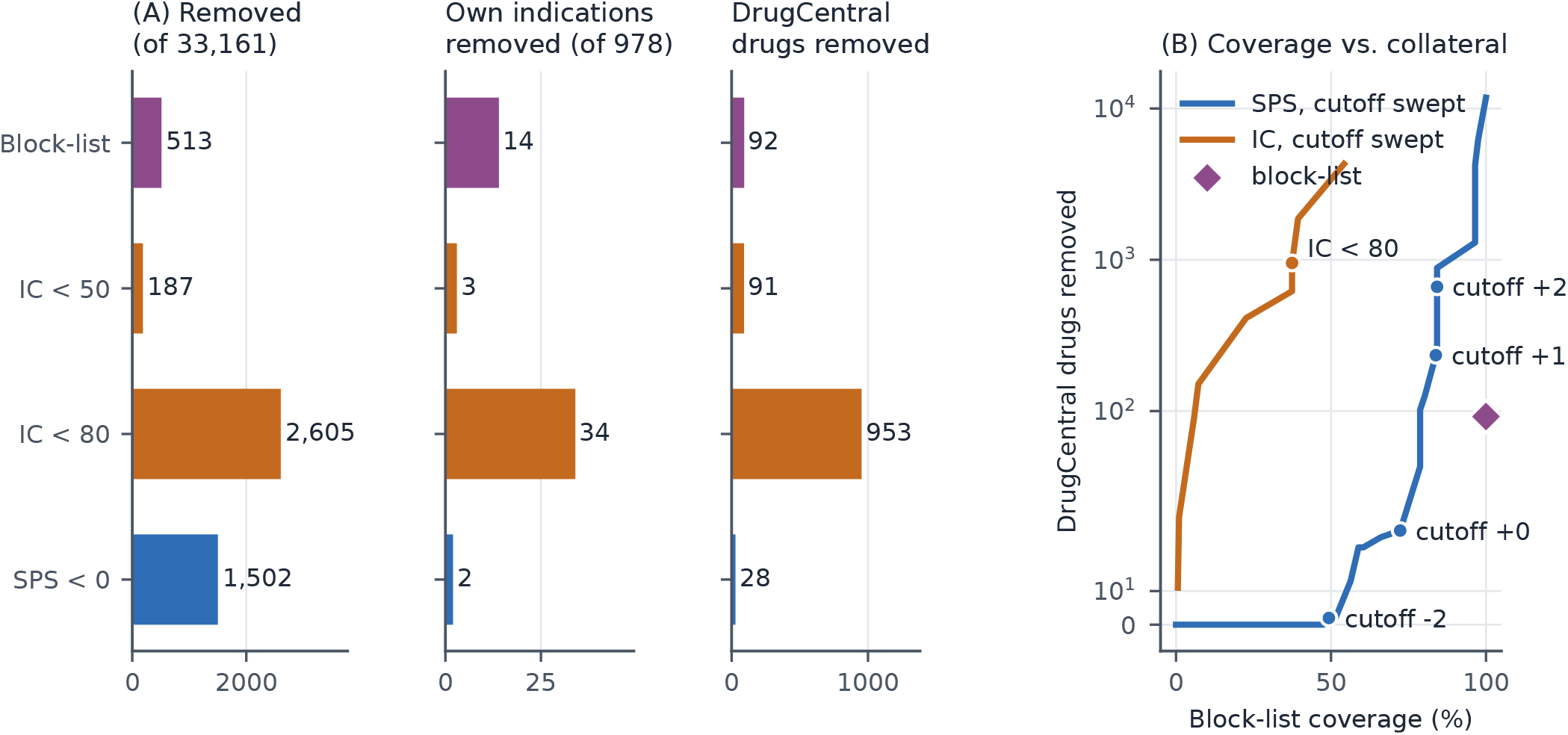
Filtering the complete answer lists of the 200 development-set `treats` queries. **(A)** Answers removed, the query disease’s own DrugCentral indications removed, and any DrugCentral drug removed, per filter (IC arms pass answers without IC). **(B)** Block-list coverage against DrugCentral drugs removed as each score’s cutoff is swept; the ARAX block-list is a single point.

In a threshold-free comparison of the two scores (SPS and IC) for filtering, the SPS filter had better recall than the IC filter at all levels of false positive error rate, as measured by the number of known-specific (DrugCentral) drugs removed (Fig. 4B). As expected for a hand-curated filter list, the ARAX generic concept block-list is conservative and precise; filtering with it led to 513 removals and only 14 known-correct answers lost, with one systematic error. However, at a threshold where the SPS filter’s false positive rate equals that of the block-list (approximately 100 DrugCentral drugs removed), the SPS filter removes 79% of the block-list concepts that appear as answers (compared to 100% with the block-list filter).

To explore the utility of the SPS for calling generic concepts beyond the ARAX generic concept block-list (which is biased toward generic concepts that frequently appear in `treats` query results), we carried out a LLM-based audit of the SPS (Sec. 2.6). On average, on List A concepts, the audit model correlated with the expert panel at *ρ* = 0.68, with the audit model over-calling generic by approximately 0.8 points on the expert-rating scale. Among the answers that the SPS removes and the block-list does not (350 sampled from 544 distinct concepts), 94% were rated generic by the audit (91 to 96) and 3% were rated specific. Among the retained answers with scores between 0 and 1, 68% were rated generic, so the SPS cutoff at 0 is conservative and some broad concepts remained among the answers with scores just above the filtering threshold.

### 3.5 Q3. Treats query findings held up on an independent labeled data set

Our empirical study of the performance of three different ARAX query result post-filters on a replication set of 200 additional diseases (Sec. 2.6) produced 91,203 result answers (a median of 155 per disease), of which 1,296 answers corresponded to approved treatments for the query disease. Of the four filters tested, the SPS filter performed best (Table 2): the block-list filter removed 1,419 answers and dropped 17 known-correct answers (known drug-disease indications); the IC*<* 50 filter removed 488 answers and dropped one known-correct answer; the IC*<* 80 filter removed 7,207 answers and dropped 58 known-correct answers; and the SPS removed 5,135 answers, covering 77% of the block-list’s removals and dropped no known-correct answers. Of all four filters, the SPS filter removed the fewest (71) approved-drug answers, and it did not reduce the number of known-positive results that were in the top-ten answers. Seventy-three percent of the concept nodes that the SPS removed had no IC.

**Table 2:** Filtering the complete answer lists of the 200 replication diseases (91,203 answers, 1,296 own-indication answers present) with the SPS model frozen. Brackets indicate 95% CI from query-level resampling. The IC filters do not remove answers that do not have an IC score. Unfiltered lists hold 837 indications in the top ten.

| Filter | Removed | Own indications removed | DrugCentral drugs removed | Block-list coverage | Top-ten indications |
| --- | --- | --- | --- | --- | --- |
| Block-list | 1,419 | 17 [10–25] | 167 [142–192] |  | 829 |
| IC < 50 | 488 | 1 [0–3] | 172 [133–211] | 5% | 837 |
| IC < 80 | 7,207 | 58 [42–76] | 2,099 [1,767–2,453] | 39% | 808 |
| SPS < 0 | 5,135 | 0 [0–0] | 71 [50–96] | 77% | 838 |

## 4 Discussion

In **Q1**, we asked how well IC tracks expert judgments of concept specificity. From our results, IC is, at best, an imperfect (and for many concepts, unavailable) specificity measure. IC is by definition sensitive to a concept’s position within an ontology, but it does not meaningfully distinguish the specificities of concepts with similar IC values. Further, IC is confounded by the degree of granularity (sparsity or density) of the ontology coverage of a concept’s semantic neighborhood.

In **Q2**, we asked if we can do better than IC with a text-based SPS model that does not depend on a concept’s knowledge graph neighborhood. The SPS effectively distinguishes between broad and specific concepts even when the concepts occupy similar positions within their respective ontologies (as illustrated by correlation with expert panel ratings, within IC bands).

In **Q3**, we asked how the SPS performs, relative to IC or the block-list, for filtering results of treats-type queries. Increasing the aggressiveness of IC filtering removes more answers, but also removes more established drugs and known indications. The SPS filter removed a larger set of broad class-level answers, while preserving nearly *all known DrugCentral indications*. This suggests that the semantic-based metric is a better signal of genericness for the downstream task of filtering drug-repurposing results. Use of a concept block-list avoids some of the problems seen with aggressive IC-based filtering, but requires manually identifying generic concepts; such human-curation would be difficult to conduct at scale. The block-list also contains concepts that may not be considered generic, but that occur so frequently as predicted answers for disease-specific queries that they are classified as ‘generic answers’. One such example, from ARAX query results, is dexamethasone: because it occurs so frequently in query answers for predicted treatments, dexamethasone is included in the ARAX block-list despite being a specific drug, and was therefore removed from several queries for which it is a known indication. This demonstrates an important distinction between genericness and relevance. A concept can be specific, but nonetheless unhelpful as a query answer. The SPS is intended to measure specificity, and could therefore be used alongside a curated list that removes specific-but-unhelpful concepts for other reasons.

For the query-level performance studies of the concept filters (Q3), because the initial 200-disease set had been used during model selection, we performed a replication analysis using the frozen SPS model. In that analysis, the SPS filter removed substantially more answers than the hand-curated block-list, while preserving the known DrugCentral indications. The removal rates of DrugCentral drugs from query answers were seemingly unaffected by whether the drug concept was included in the training or not. This suggests that the SPS’s query-level filtering performance is not simply attributable to the development diseases or to the memorization of DrugCentral drugs.

We note several limitations of this work. The expert panel consisted of only three raters and 150 concepts, and the no-IC concepts had relatively low inter-rater agreement. The query-level evaluation also does not provide expert ground truth for every removed answer. DrugCentral is useful for identifying cases where a known treatment is incorrectly removed, but it cannot determine whether the remaining removed concepts are truly generic. In addition, the initial 200-disease query set was used during model selection, and DrugCentral both contributed training examples and was used in benchmarking. We addressed these dependencies with a second 200-disease replication set evaluated after the model and cutoff were frozen, and by comparing removals between DrugCentral drugs used for training and those held out from training. Finally, these results are specific to one Translator knowledge graph build and one reasoning agent, so additional work will be needed to determine how well the approach generalizes to other systems.

Overall, the results from the studies presented here support treating concept specificity as a signal that is best captured with a text-based model rather than relying solely on a block-list or ontology position. Although we evaluated query-level performance in the context of therapeutic drug recommendations, diverse applications of computational reasoning could benefit from improved answer specificity, including explaining gene–phenotype associations and analyzing genomics data.

## Author Contributions

FH conceived the study, wrote the code, and performed the data analysis. FH and SAR authored the manuscript. DK, JCR, and SAR served as the expert annotators and contributed to design, interpretation, and revision of the manuscript. SAR supervised the work. AB helped to revise the manuscript. All authors contributed to the article and approved the submitted version.

## Acknowledgments

The authors thank the ARAX team and the NCATS Biomedical Data Translator consortium members for the knowledge graph, tooling, and test infrastructure that this work builds on.

## Generative AI statement

The authors declare that generative AI was used in the preparation of this manuscript. Claude Fable 5 (Anthropic) and GPT-5.6 Sol (OpenAI) were used to draft and revise portions of the text, write and review analysis code, and verify reported values against the underlying results. All AI-generated output was reviewed and verified by the authors, who take full responsibility for the content.

Separately, locally hosted open-weight language models were used as part of the research methodology: Qwen2.5-32B-Instruct (Alibaba Cloud) was used for label review, and Gemma-3-12B-it (Google), Mistral-Nemo-Instruct-2407 (Mistral AI), and Phi-4 (Microsoft) were used as audit judges.

## Data Availability Statement

The datasets generated and analyzed for this study, including panel ratings, concept list definitions, trained models, and the responses for the three 200-disease query runs, are available from the corresponding author. The code for reproducing the analyses and figures is available at github.com/hodgesf/semantic-score.

