## supplemental information for "Validated Semantic Scoring Improves Filtering of Generic Concepts from Knowledge Graphs for Drug Repurposing Queries"

### Supplementary Information

#### S1 Expert-Set Construction Details

Concepts used as labeled examples in Semantic Precision Score (SPS) training were excluded from both expert-rated lists. List B additionally excluded concepts in earlier curated review files, the first-stage training set, and List A. Both lists were fixed before annotation.

To limit near duplicates, concept names were lower-cased and split into alphanumeric tokens. A candidate was rejected when its token set contained, or was contained by, that of a previously selected concept. For List B, this check also included List A. Candidates were considered in a seeded random order, so the surviving member of a near-duplicate group depended on that order.

Within each IC band, List A candidates were accepted subject to a cap of three per Biolink category. One band required a cap of four to reach its target of 20 concepts; the near-duplicate rule remained in force. List B used fixed category quotas: Drug (8), ChemicalEntity (7), SmallMolecule (5), Disease (5), Pathway (5), GeneFamily (5), PhenotypicFeature (4), Procedure (4), Protein (3), MolecularMixture (2), and AnatomicalEntity (2). Concepts were sampled randomly within each category. List A was drawn under seed 2654 and List B under seed 2655

#### S2 First-Stage Genericness Model

##### S2.1 Construction of Parent–Child Training Pairs

Pairs were inferred from concept names and definitions without using concept ontology edges, IC values, or expert ratings. For lexical pairs, names were normalized by lower-casing, un-inverting MeSH-style headings, removing trailing parenthetical qualifiers, and tokenizing and singularizing the final token. A candidate parent was accepted when its normalized name was a proper suffix of the child's name; all matching suffix lengths were considered.

Lexical matches were rejected when the token before the match indicated a relational or negated phrase, such as "of" in "regulation of transport" or "non" in "non small cell lung carcinoma." Matches involving short codes or numeric fragments were also rejected. For both lexical and definition-derived pairs, the concepts had to have compatible Biolink categories, meaning that either category occurred in the other's category lineage.

For definition-derived pairs, the genus phrase was extracted from the first sentence of an OBO-style definition and matched to another concept by normalized name, with lexical suffix matching as a fallback. Pairs that resolved to the concept itself were discarded. After deduplication, the two rules yielded 660,663 distinct child–parent pairs.

#### S2.2 Genericness-Score Estimation

Concept text was encoded with `cambridgeltl/SapBERT-from-PubMedBERT-fulltext`, using "name: description" when a description was available and the name alone otherwise. We used the final-layer CLS representation, truncated inputs to 128 word pieces, and $L_2$-normalized each embedding $\mathbf{e}_x \in \mathbb{R}^{768}$.

The 660,663 parent–child pairs were shuffled with seed 2654, then capped sequentially at 30 pairs per parent and 150,000 pairs per child's Biolink category. This left 332,184 pairs and limited the influence of unusually broad parents and high-volume categories.

For each retained child–parent pair $(c, p)$, we formed $\mathbf{d}_{cp} = \mathbf{e}_p - \mathbf{e}_c$. An intercept-free logistic classifier received $\mathbf{d}_{cp}$ with label 1 and $-\mathbf{d}_{cp}$ with label 0. Its modeled probability of the parent-first ordering is $\sigma(\mathbf{w}^\top \mathbf{d}_{cp})$, where $\sigma$ is the logistic function. Reversing the pair negates the logit, and both orientations contribute the same loss. The resulting per-pair objective is

$$\mathcal{L}(\mathbf{w}) = \frac{1}{|\mathcal{P}|} \sum_{(c,p)\in\mathcal{P}} \log\bigl(1 + \exp[-\mathbf{w}^\top(\mathbf{e}_p - \mathbf{e}_c)]\bigr) + \frac{\alpha}{2}\|\mathbf{w}\|_2^2, \qquad \alpha = 10^{-5}$$

where $\mathcal{P}$ is the retained pair set. We fitted the classifier with stochastic gradient descent for 30 passes, using random state 2654 and disabling convergence-based stopping.

Because $\mathbf{w}^\top(\mathbf{e}_p - \mathbf{e}_c) = r(p) - r(c)$ for $r(x) = \mathbf{w}^\top \mathbf{e}_x$, the fitted coefficients also score individual nodes. We standardized these projections across all graph nodes as

$$g(x) = \frac{r(x) - \mu_r}{\sigma_r}$$

where $\mu_r$ and $\sigma_r$ are the graph-wide mean and standard deviation. Higher $g(x)$ indicates greater genericness. For comparisons with IC, we plotted $-g(x)$ so that both plotted scores increase with specificity.

#### S2.3 Validation of the Genericness Score

After pair balancing, we randomly held out 20% of distinct parents and fitted the model on pairs with other parents. On the 67,039 held-out pairs, the parent scored above the child in 91.9% of cases. We then discarded this fit and refitted the model on all 332,184 balanced pairs; the resulting score was used in subsequent analyses.

For each AUC analysis, comparison concepts were sampled without replacement within Biolink categories, up to 20 per curated generic concept in that category. Curated and oracle-confirmed generic concepts were excluded from the comparison pool. The pair-ablation and production-score analyses used separate draws of this pool.

To assess stability, we fitted separate models on two disjoint halves of the parents. Their scores had Spearman correlation $\rho = 0.987$ across 300,000 randomly sampled graph nodes. Models fitted using only lexical or only definition-derived pairs achieved ROC-AUCs of 0.907 and 0.932, respectively, against 0.926 for an all-pairs model fitted and evaluated in the same way. The lexical-only and definition-only scores differed substantially in their node rankings (Spearman $\rho = 0.476$).

The refitted genericness score achieved ROC-AUC 0.927 against its sampled comparison pool, versus 0.778 for node degree. Within five equal-sized strata ordered by $\log(1 + \text{degree})$, score ROC-AUC ranged from 0.889 to 0.954, including 0.954 in the lowest-degree stratum. Because comparison concepts were unlabeled, these AUCs measure separation from the sampled pool rather than accuracy against confirmed specific concepts.

### S3 Positive-Unlabeled Candidate Discovery

#### S3.1 Initial Labels and Pseudo-Negative Sampling

The 4,193 identifiers in the production ARAX block-list were normalized through Babel and matched to preferred or equivalent identifiers in the tier-0 graph, yielding 1,053 initial positive nodes.

At each training round, we drew $10|P_t|$ nodes without replacement from the unlabeled pool, using seed 2654, and treated them as pseudo-negatives. The draw was uniform across categories, degree, IC availability, and classifier features. The pool and draw were updated after each review round to exclude newly confirmed positives. Because the block-list is incomplete, pseudo-negatives could still include generic concepts.

Oracle-confirmed specific concepts were retained as hard negatives and added to the sampled pseudo-negatives in subsequent rounds. Thus, the 10:1 ratio describes the initial draw, rather than the final balance after filtering and adding hard negatives.

#### S3.2 GBDT Features

The positive-unlabeled candidate classifier used the ten features in Table S1. Counts use incident edges unless the definition specifies distinct neighbors. Predicted IC for nodes with observed IC was generated out of fold, so a node's IC was never used to fit its own prediction. Ridge penalties were selected from $0.001, 0.01, 0.1, 1, 10, 100$ using the nested fold procedure with seed 2654; nodes without observed IC were scored by a model fitted to all observed-IC nodes at the modal selected penalty. Features were neither standardized nor imputed. Missing values were passed directly to XGBoost.

**Table S1.** Features used by the candidate classifier.

| Feature | Definition |
|---|---|
| `degree` | Number of incident edges, including multiple predicates between the same nodes. |
| `unique_neighbors` | Number of distinct adjacent nodes. |
| `distinct_neighbor_cats` | Number of Biolink categories among adjacent nodes. |
| `neighbor_cat_entropy` | Shannon entropy of adjacent-node categories, counted per incident edge. |
| `predicate_entropy` | Shannon entropy of predicates on incident edges. |
| `hierarchical_child_count` | Number of incoming `biolink:subclass_of` edges, including duplicates. |
| `information_content` | Observed tier-0 IC; missing where unavailable. |
| `predicted_ic` | Ridge-regression estimate of IC from the SapBERT embedding and most specific Biolink category. |
| `ic_minus_predicted_ic` | Observed IC minus predicted IC; missing where observed IC is unavailable. |
| `cos_to_category_centroid` | Cosine similarity to the mean SapBERT embedding of nodes in the same Biolink category. |

We excluded the node's Biolink category as a direct classifier feature after development runs showed strong category-level predictions that could concentrate subsequent oracle review within particular categories. This was a development decision, not a controlled ablation; no paired runs were retained. Category still enters indirectly through predicted IC, similarity to the category centroid, and the categories of neighboring nodes. All four reported training rounds used the final feature set without direct category.

#### S3.3 GBDT Training and Iterative Review

At round $t$, let $P_t$ contain the block-list seed nodes and oracle-confirmed generic concepts ($|P_0| =$ 1,053), and let $H_t$ contain oracle-confirmed specific concepts ($H_0 =$). A fresh pseudo-negative sample $N_t$ was drawn from nodes in neither set, with $|N_t| = 10|P_t|$. The training set was $T_t = P_t \cup H_t \cup N_t$, so

$$|T_t| = 11|P_t| + |H_t|, \qquad \texttt{scale\_pos\_weight} = \frac{10|P_t| + |H_t|}{|P_t|}.$$

Hard negatives were therefore added to, rather than substituted for, the 10:1 pseudo-negative draw.

The XGBoost model used 300 trees, maximum depth 4, learning rate 0.05, row and column subsampling of 0.8, and the histogram tree method. Within each round, we measured out-of-fold average precision using five-fold stratified cross-validation (seed 2654). The positive fraction ranged from 0.078 to 0.091 across rounds, providing an approximate random-ranking baseline.

After fitting, the model scored all 1,754,754 graph nodes. Unlabeled nodes scoring above 0.8 were submitted for oracle review; the threshold itself assigned no labels. Reviewed generic and specific concepts were added to form $P_{t+1}$ and $H_{t+1}$ before the next draw and fit. The round 3 queue was not reviewed.

**Table S2.** Candidate-discovery rounds. Average precision was measured out of fold within each round. 'Above 0.8" is the number of unlabeled nodes queued for review; 'Oracle G / S" gives the numbers confirmed generic and specific. The round 3 queue was not reviewed.

| Round | Training rows | Positives | Negatives | Average precision | Above 0.8 | Oracle G / S |
|---|---|---|---|---|---|---|
| 0 | 11,583 | 1,053 | 10,530 | 0.9609 | 21,689 | 7,058 / 14,631 |
| 1 | 103,852 | 8,111 | 95,741 | 0.6907 | 2,747 | 1,384 / 1,363 |
| 2 | 120,439 | 9,495 | 110,944 | 0.6934 | 1,159 | 538 / 621 |
| 3 | 126,978 | 10,033 | 116,945 | 0.6912 | 957 | — |

Average precision describes a different training mixture in each round: later rounds contain specific concepts that earlier models scored highly and sent for review. The queue shrank from 21,689 to 957 across the four fits. We stopped for practical reasons, not because formal convergence had been established.

### S4 LLM Oracle Implementation

Candidate concepts were reviewed with `Qwen/Qwen2.5-32B-Instruct-AWQ`, served locally through vLLM with temperature 0. Each request contained ten numbered concepts. For each concept, the oracle received its name and Biolink category, but no identifier, GBDT score, IC value, graph statistics, or earlier verdict. The category was omitted for `biolink:GeneFamily` nodes because that source category also contains individual entities, for which the "family" label could bias the verdict.

Decoding was constrained to a JSON schema. A batch was accepted only if it returned exactly one `G` or `S` verdict for each numbered concept; otherwise, the entire batch was retried. All submitted concepts received a valid verdict.

### S5 Final Semantic Score Training Set

The Semantic Precision Score (SPS) training set was assembled separately from the PU classifier's pseudo-negative samples. Positives came from the curated block-list and oracle-confirmed generic concepts. Manual review removed 28 erroneous generic labels, chiefly gene and chemical records. Sample A concepts were excluded from the assembled pools.

Negatives were assembled only within Biolink categories containing positives, targeting three per positive. Within each category, eligible oracle-confirmed specific concepts were used first, up to that target; any remainder was filled by sampling unlabeled concepts from the same category. Oracle-confirmed specifics from `AnatomicalEntity`, `GrossAnatomicalStructure`, `Cell`, and `CellularComponent` were excluded from the hard-negative pool. This exclusion did not apply to unlabeled top-up draws.

The subjects of DrugCentral `biolink:treats` edges were split by the MD5 hash of their identifiers, with even hashes assigned to the training half. Held-back-half drugs were removed from the assembled set, including any already present. Next, 3,188 negative rows whose node identifiers began with `PUBCHEM.COMPOUND` or `CHEMBL.COMPOUND` were removed without replacement. Training-half DrugCentral drugs not already present were then added as negatives. The held-back half was used for epoch selection together with block-listed query answers.

Before the internal holdout, the resulting set contained 10,004 positives (1,051 curated; 8,953 oracle-confirmed) and 24,911 negatives (9,847 oracle-confirmed; 14,045 sampled unlabeled; 1,019 DrugCentral). After shuffling with seed 2654, the first 10% of rows were held out to report validation AUC after each epoch. They were not used to select the epoch.

##### S5.1 DrugCentral Split

DrugCentral examples were the 2,330 distinct subjects of `biolink:treats` edges whose provenance included `infores:drugcentral`. An identifier was allocated to training when its MD5 digest, interpreted as an integer, was even, and held back otherwise. This allocated 1,127 drugs to training and 1,203 to the held-back set. After the training-set exclusion rules, 1,019 of the allocated drugs remained as negative examples.

The held-back drugs were used for epoch selection (Section S6.1). They therefore provide a check on drugs absent from parameter fitting, but not a fully independent test of the selected model.

#### S6 Semantic Score Model: Implementation Details

We used `FremyCompany/BioLORD-2023` with inputs formatted as "name: description," or the name alone when no description was available, truncated to 96 word pieces. The attention-mask-weighted mean of the final-layer token representations was $L_2$-normalized before concatenation with the standardized first-stage genericness score. Generic concepts had label 1. The binary cross-entropy loss weighted that class by $N_-/N_+$, calculated on each run's training split (approximately 2.49). Scores were treated as discriminative logits; no probability calibration was performed.

Optimization used AdamW with weight decay 0.01, bfloat16 mixed precision, and gradient-norm clipping at 1.0. The encoder and classification head used learning rates of $2 \times 10^{-5}$ and $10^{-3}$, respectively. Runs used seeds 2654, 2, and 3 and trained for at most three epochs. The reported SPS is the negative of the mean of the three selected logits, rather than the mean of their probabilities.

#### S6.1 Epoch Selection

For each seed, we selected an epoch without using expert-panel ratings. Among epochs that classified at most 1% of the 1,203 held-back DrugCentral drugs as generic ($S < 0$), we chose the one maximizing the difference between the proportion of development-query block-list removals classified as generic and the proportion of held-back drugs classified as generic. If no epoch met the 1% constraint, we retained the first epoch.

Because the first 200 disease queries and held-back DrugCentral drugs informed epoch selection, results on those data are development results. The second 200-disease replication set was evaluated after the seed-specific epochs, ensemble rule, and zero cutoff were fixed.

### S7 ARAX Query Implementation

Queries used RTX commit `2d1fb2d1` and tier-0 build 2026-06-21 with empty caches. Generic-node removal and the 500-result cap were disabled during collection. Each filter was then applied offline to the same saved, ranked TRAPI response, followed by the production cap of 500 results. An answer was the node bound to the treatment slot.

The block-list arm used ARAX's production matching rules for identifiers, equivalent identifiers, names, and its regular-expression pattern. Eligible diseases were MONDO-identified `biolink:Disease` nodes with at least two distinct DrugCentral indications; 48 Translator sprint-6 test diseases were excluded. We sampled 200 of the remaining 687 diseases with seed 2654.

Reported counts sum *(query, answer)* pairs, so the same drug removed from several queries is counted several times. Retriever timed out for 75 of the 200 development queries. Because all filters operated on each disease's same saved response, a timeout did not favor one filter arm over another, but it reduced the available query results.

#### S7.1 Semantic Score Threshold Sweep

Development-query answers were filtered at the cutoffs in Table S3; an answer was removed when its SPS was below the cutoff. The 0.46 cutoff was identified on the development set because it matched the production block-list's 92 DrugCentral answer removals. It is a descriptive, matched operating point, not a separately selected deployment threshold.

**Table S3.** Threshold sweep on the development queries. Counts are (query, answer) pairs; coverage is the percentage of the block-list's 513 removals also removed at that cutoff.

| Cutoff | Answers removed | DrugCentral removals | Own indications removed | Block-list coverage |
|---|---|---|---|---|
| −1.00 | 1,222 | 13 | 0 | 56.3% |
| 0.00 | 1,502 | 28 | 2 | 72.3% |
| 0.46 | 1,889 | 92 | 4 | 78.8% |
| 1.00 | 2,289 | 234 | 7 | 83.8% |

### S8 LLM Audit

We screened three locally hosted, 4-bit open-weight judges against expert-rated List A before auditing query answers. Gemma-3-12B-it and Phi-4 exceeded the prespecified Spearman correlation threshold of 0.50 ($\rho = 0.62$ and 0.75); Mistral-Nemo-Instruct-2407 did not ($\rho = 0.46$). The two retained judges were averaged. These correlations describe the model-selection set, not independent validation of the judges.

The retained judges rated all 150 expert-list concepts and 620 concepts sampled from query answers. Query concepts were sampled across filter-agreement groups, with further stratification by SPS range and IC availability. This deliberately emphasized disagreements between filters; the resulting ratings do not estimate genericness prevalence among all answers. Judges were blinded to IC, SPSs, block-list membership, and expert ratings. The prompt and sampling rules were fixed before query-answer ratings were inspected; the prompt and sampled identifiers are provided with the analysis code.

### S9 Replication and DrugCentral Split Analysis

After the models, selected epochs, ensemble rule, and zero cutoff were fixed, we sampled 200 replication diseases from the 687-disease eligibility pool, excluding the 200 development diseases (seed 2655). Retriever contributed results to 191 replication queries.

Among DrugCentral answer occurrences in the replication responses, the SPS removed 50 of 17,735 (0.28%) involving drugs allocated to the training half and 21 of 18,009 (0.12%) involving held-back drugs. At the distinct-drug level, 3 of 1,016 training-half drugs and 4 of 1,083 held-back drugs scored below zero. These are descriptive comparisons; the held-back drugs also informed epoch selection.

### S10 Statistical Details

Spearman correlations used midranks for ties, and Krippendorff's $\alpha$ used the ordinal disagreement metric for the five-point ratings. Concept-level 95% confidence intervals were percentile intervals from 10,000 bootstrap resamples. When comparing two correlations on the same concepts, both were recalculated on each shared resample before taking their difference.

For replication-query confidence intervals, 2,000 bootstrap resamples drew disease queries with their complete answer lists, preserving clustering of answers within a query.

### S11 Software and Reproducibility

Analyses used Python 3.12 in separate CPU and GPU environments. Candidate discovery used XGBoost 3.3.0; model training and local inference used PyTorch 2.11.0, Transformers 5.14.1, and vLLM 0.26.0 under CUDA 13. GPU work ran on a single NVIDIA RTX PRO 4000 Blackwell with 24 GB of memory. The analysis repository cited in the Data Availability Statement contains the code and materials needed to reproduce the reported results.

### Figures

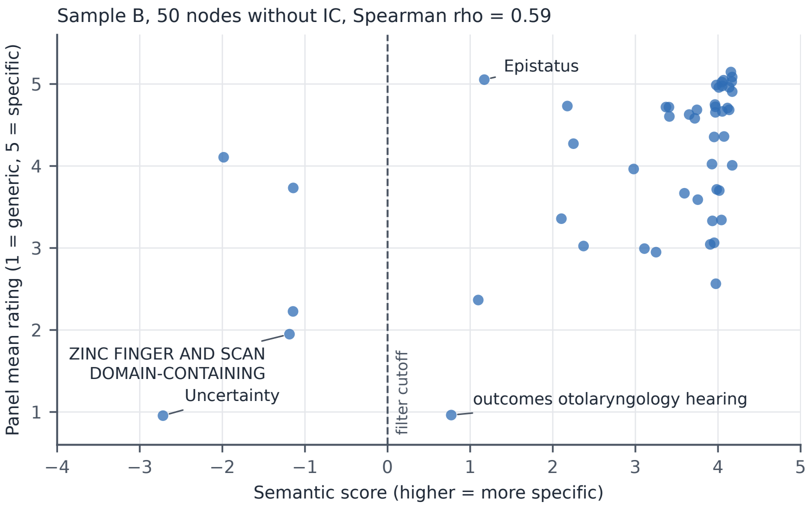


Figure S1: Semantic Score against the panel mean on List B (50 concepts without IC), Spearman $\rho = 0.59$. Panel means are averages of three integer ratings and are jittered vertically for legibility; scores are unmodified. The dashed line is the filter cutoff $S = 0$. Labelled points are the three concepts the panel rated generic, and Epistatus, the largest disagreement in the opposite direction (panel mean 5.0, score 1.15).
